# 8-HQA Adjusts the Number and Diversity of Bacteria in the Gut Microbiome of *Spodoptera littoralis*

**DOI:** 10.1101/2022.12.06.519313

**Authors:** Tilottama Mazumdar, Sabine Hänniger, Shantanu P. Shukla, Aishwarya Murali, Stefan Bartram, David G. Heckel, Wilhelm Boland

## Abstract

Quinolinic carboxylic acids are known for their metal ion chelating properties in insects, plants and bacteria. The larval stages of the lepidopteran pest, *Spodoptera littoralis*, produce 8-hydroxyquinoline-2-carboxylic acid (8-HQA) in high concentrations from tryptophan in the diet. At the same time, the larval midgut is known to harbor a bacterial population. The motivation behind the work was to investigate whether 8-HQA is controlling the bacterial community in the gut by regulating the concentration of metal ions. Knocking out the gene for kynurenine 3-monooxygenase (KMO) in the insect using CRISPR/Cas9 eliminated production of 8-HQA and significantly increased bacterial numbers and diversity in the larval midgut. Adding 8-HQA to the diet of knockout larvae caused a dose-dependent reduction of bacterial numbers with minimal effects on diversity. *Enterococcus mundtii* dominates the community in all treatments, probably due to its highly efficient iron uptake system and production of the colicin, mundticin. Thus host factors and bacterial properties interact to determine patterns of diversity and abundance in the insect midgut.

## 1 Introduction

All higher animals harbor a dynamic community of microorganisms that they have acquired along their life-cycle, the population of which they control in a way that the pathogens are eliminated. Such is the proximity between these two entities that the gut bacteria act as a major organ, influencing the health and fitness of their host. This community is continuously shaped along the life cycle of the organism, jointly by its diet, behavior, physicochemical environment of the gut, and host-genetics. The colonization of microbes in the gut begins at birth by vertical transmission, (Blekhman et al., 2015) and then it is largely shaped by physicochemical condition of the host gut (Engel and Moran, 2013). From the point of view of arthropods, guts of lepidopterans have an alkaline pH, owing to a tannin-rich diet, hence rejecting several bacteria (Berenbaum, 1980). Some insects employ their innate immune pathways for regulating bacterial populations, for example green weevils and their anti microbial peptides (AMP) (Login et al., 2011). Formicine ants swallow their acidic antimicrobial gland secretion to keep opportunistic pathogens in check (Tragust et al., 2020).

*Spodoptera littoralis* or the cotton leaf worm is an agricultural pest dominant in tropical and sub-tropical areas. It exhibits a holometabolous mode of development with distinct larval, pupal and adult stages. The gut microbiota of *S. littoralis* has been elucidated. In its larval stages, clostridia and enterococci constitute most of the population, followed by enterobacteriaceae, although a dynamic trend of bacterial composition was observed over the course of its development. *Enterococcus mundtii* starts dominating from the second instar onwards and persists throughout the larval period (Tang et al., 2012). Certain important factors influencing the dynamic population in their guts is an extensive metamorphosis owing to their holometabolous life cycle, a pH gradient of alkaline to neutral along the length of the gut, and presence of an iron chelator, 8-hydroxyquinoline-2-carboxylic acid (8-HQA, 8-hydroxyquinaldic acid) produced by the larvae in their gut regurgitant. This compound is synthesized by the host from dietary tryptophan and is present in high concentration: 0.5-5 mM in the regurgitant of the larvae (Pesek et al., 2015). Quinolinic carboxylic acid derivatives are widely distributed in Nature. One of their uses is quorum sensing in bacteria. 8-Hydroxy-4-methoxyquinoline-2-carboxylic acid is a siderophore, used for high-affinity uptake of iron, in *Pseudomonas fluorescens* (Mossialos et al., 2000). Quinolinic carboxylic acids have also been found in plants (Schennen and Hölzl, 1986). 8-HQA additionally acts as a chelator for divalent metal ions. Electron Spin Resonance (ESR) measurements have demonstrated that the affinity of 8-HQA towards Fe^3+^ is higher than that of Fe^2+^ (Gama et al., 2018).

Since such compounds have been proven to have siderophoric roles and affect the availability of metal ions in the gut, we hypothesized that 8-HQA might regulate the gut bacterial population as well. Among all metal ions, iron especially is an important microelement since it takes part in diverse cellular processes like nucleic acid synthesis, tricarboxylic acid cycle, electron transport, secondary metabolism and functions as a cofactor in proteins and enzymes (Andrews et al., 2003). Hence, gut bacteria should come up with efficient methods to take up iron from their surroundings.

To test whether 8-HQA controls the bacterial population in the gut of *S. littoralis*, the genes for enzymes in the insect’s biosynthetic pathway were knocked out using CRISPR/Cas9. The well-known pathway from tryptophan to xanthurenic acid (Han et al., 2007) is shown in Figure 1. Although the final steps producing 8-HQA are unknown, deuterium labelling experiments have shown that 3-hydroxykynurenine is a precursor (Pesek et al., 2015), so we knocked out kynurenine 3-monooxygenase (KMO) which converts kynurenine to 3-hydroxykynurenine, and two putative transaminases (AGT1 and AGT2) converting 3-hydroxykynurenine to xanthurenic acid. No viable AGT1 knockouts were recovered, and AGT2 knockouts produced normal levels of 8-HQA (data not shown). However, KMO knockouts were viable, and no 8-HQA could be detected in their regurgitant. Comparison of the bacterial communities of wild-type and knockout larvae, with or without external supplementation of 8-HQA, showed that 8-HQA has significant effects on bacterial abundance and diversity in the midgut of *S. littoralis* larvae.

**Figure 1.**
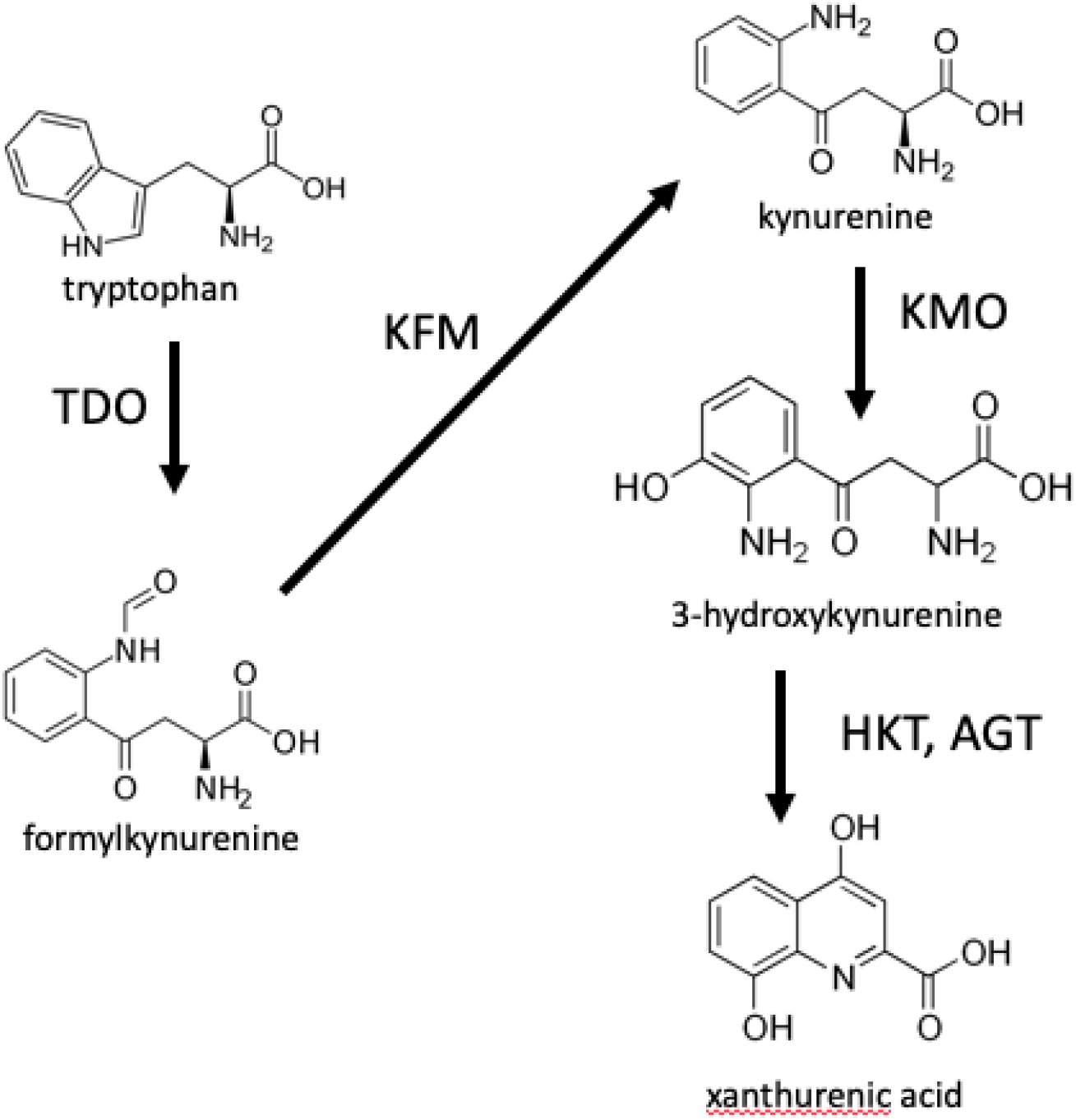
Initial steps in the biosynthesis of 8-HQA. Tryptophan is converted to formylkynurenine by tryptophan dioxygenase, which is then converted to kyurenine by kynurenine formamidase. Kynurenine in turn is converted to 3-hydroxykynurenine by kynurenine monooxygenase (KMO). The toxic 3-hydroxykynurenine is converted to xanthurenic acid by transamination (here shown via alanine glyoxalate aminotransferase, represented by two enzymes AGT1 and AGT2 encoded by separate genes in *S. littoralis*) followed by cyclization. Labelling experiments have shown that 3-hydroxykynurenine is a precursor of 8-HQA (Pesek et al., 2015) therefore enzymes involved in its synthesis and conversion were targeted by CRISPR/Cas9.

## 2 Materials and Methods

### 2.1 Insect Rearing and Sampling

*S. littoralis* were obtained from an old laboratory strain of the Institut National de la Recherche Agronomique, Versailles, France, which had been maintained on a semi-artificial diet since the 2000s on a semi-artificial diet (Durand et al., 2010;Meslin et al., 2022). Knockout of the gene encoding kynurenine 3-monooxygenase was carried out by the CRISPR/Cas9 method as described below. KMO-ko larvae (homozygous for the knockout) and WT larvae (wild type) were grown in separate petri dishes containing white lima bean based artificial diet as described by (Spiteller et al., 2000). This food was mixed with the frass (frass: food = 1:5 w/w) from larvae of a different laboratory strain purchased from Syngenta Crop Protection Münchwilen AG (Münchwilen, Switzerland), because the previous experiments that studied the bacterial population of *S. littoralis* larvae were conducted on this laboratory strain (Tang et al., 2012;Teh et al., 2016). Hence, the microbial population of the Syngenta line was allowed to colonize the Versailles line (WT and KMO-ko), to maintain uniformity in the gut population among these two strains. The larvae were separately reared and maintained based on family numbers to prevent them from inbreeding.

Five larvae from second, third, fourth and fifth instar stages of the KMO-ko mutant and WT lines fed with artificial diet spiked with frass from Syngenta strain, were sampled and collected. The insects were frozen for 20 min before the dissection, after which they were surface sterilized in 70% ethanol for three times, followed by rinsing in sterile water. All the samples were flash frozen in liquid nitrogen and stored at -80°C until DNA extraction. Three technical replicates containing 0.25 g of food mixed with the frass was also sampled the same way.

8-Hydroxyquinoline-2-carboxylic acid (98%, ACROS Organics) was dissolved in distilled water with pH adjusted to 10 using a solution of 1 M sodium hydroxide for better solubility. This way, a gradient of concentration of 8-HQA was fed to batches of KMO larvae: 1 mM, 3 mM, 5 mM and 10 mM: 0.8-1 g diet, plus 100 μl of the solution containing the aforementioned final concentrations of 8-HQA, in order to approximate the concentration of 8-HQA naturally occuring in WT larvae. A control batch of the same (termed as 0 mM 8-HQA) was fed with diet spiked with 100 μl of the solvent. Five individuals from each concentration were sampled in the same way.

### 2.2 Mutagenesis of 8-HQA Biosynthesis Using CRISPR/Cas9

The CRISPR/Cas9 system was used to obtain a knockout line of *S. littoralis* that does not produce 8-HQA. In total, three target genes were knocked out in *S. littoralis* embryos in this study. Initially, two genes potentially coding for alanine-glyoxylate aminotransferase, *AGT1* (GenBank Accession ON808443) and *AGT2* (GenBank Accession ON808444), were identified based on a study on mosquitoes (Rossi et al., 2005) from an in-house *S. littoralis* transcriptome. Subsequently, kynurenine 3-monooxygenase (*KMO*, GenBank Accession ON808442) was identified from the transcriptome. The ZiFit Targeter version 4.2 website (zifit.partners.org) was used to identify targets for the knockout of the three genes with the Cas9 endonuclease (target sequences in Table 1).

**Table 1:**
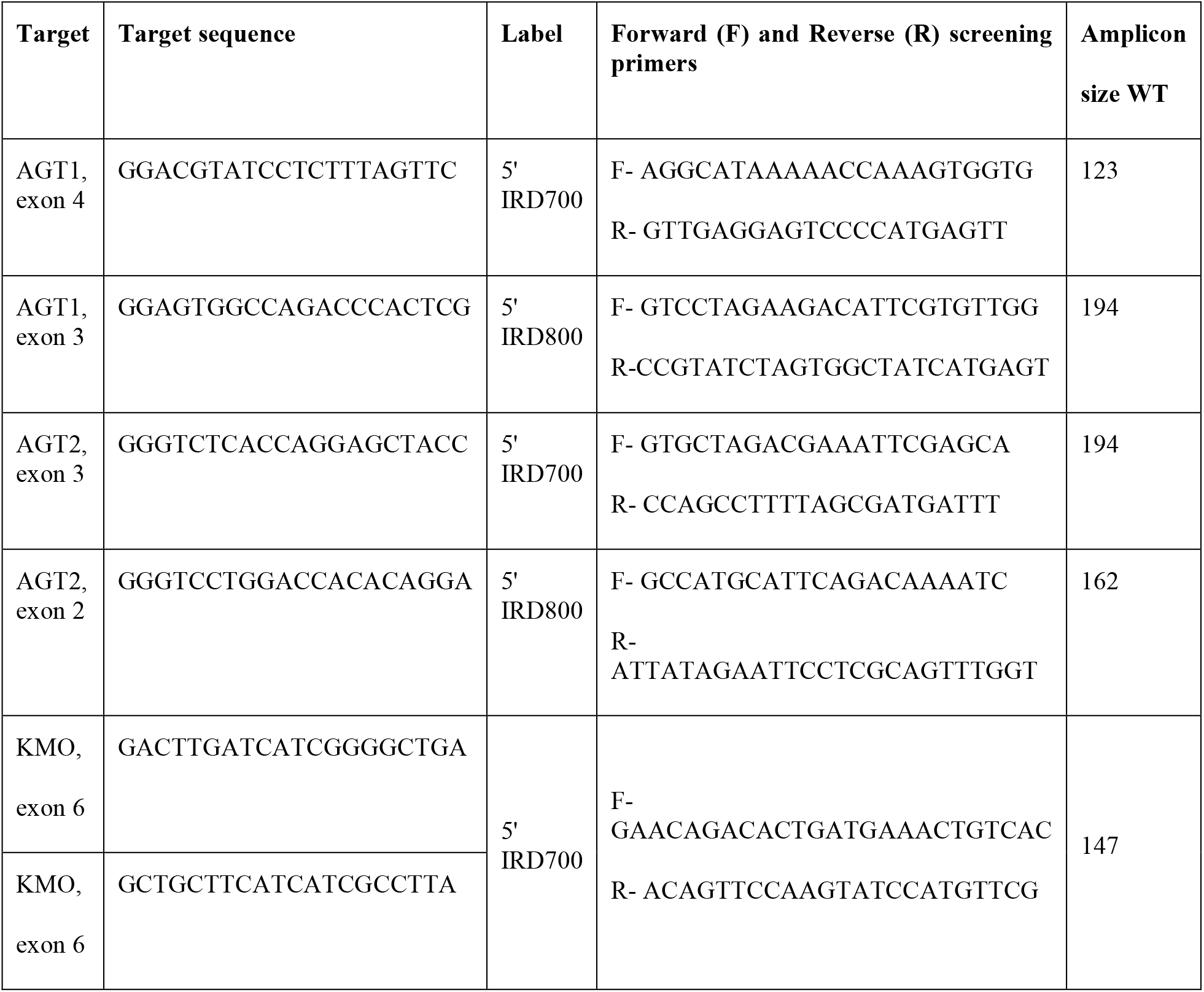
Target and primer sequences for the different knockout target genes.

In the case of *AGT1* and *AGT2*, sgRNA and Cas9 mRNA were synthesized *in vitro* and injected into freshly-laid eggs as described in Khan, Reichelt et al. (2017)). The concentration of the injection solution was 100ng/μl sgRNA and 250ng/μl Cas9 mRNA.

For *KMO*, tracrRNA, crRNA and Alt-R® S.p. Cas9 Nuclease were purchased from IDT (Integrated DNA Technologies, BVBA, Leuven, Belgium) and processed according to the manufacturer’s instructions. The injection solution contained 10 pmol tracrRNA+crRNA and 5 pmol Cas9 enzyme per 1 μl.

Single pair matings were set up in paper cups covered with gauze. Oviposition was monitored and egg masses on the gauze were collected and injected within 1h after oviposition. The scales and top layers of the egg masses were removed using sticky tape. The eggs were injected with the different injection solutions loaded into a self-made glass needle. Glass needles were made by pulling a capillary (GB100TF-10, 0.78 × 1.00 × 100 mm; Science Products, Hofheim, Germany) on a P-97 micropipette puller (Sutter Instrument, Novato, California). Tips of the needles were cut with a razor blade. Injections of egg masses were done with an Eppendorf FemtoJet 5247 at 50 hPa resting pressure, 500 hPa injection pressure. Injected egg masses were held in a humidified chamber at room temperature until hatching. Surviving larvae were screened for mutations and reared to adulthood.

For mutation screening, DNA was extracted from a single larval leg, following (Haenniger et al., 2020) with the modification that only 200 μl of the Chelex® 100 Resin solution was used per sample. Each target region was amplified with PCRs with 5% of the forward primer substituted with a 5’IRD700 or 5’IRD800 labelled primer (Table 1). This way the amplicon could be detected in a Licor 4300 DNA analyzer (LI-COR Biosciences, Bad Homburg v.d.H., Germany). 96 samples were run simultaneously and the visual output allowed for detection of insertions and deletions as small as 2 bp and up to a size of ∼1000 bp.

No *AGT1* injected eggs survived to hatching, possibly due to accumulation of the toxic 3-hydroxykynurenine (Figure 1). For *AGT2*, one surviving larva which developed into a female showed 2 different mutations in the germline, producing two different mutant lines: one with a 2bp insertion causing a frame shift in the gene, and one with a 3 bp insertion (Figure 2). Subsequent regurgitant measurements in these mutant lines showed 8-HQA levels similar to the WT larvae (data not shown). Thus, the *AGT2* line was discontinued and *KMO* was used as the new target gene for knockout.

**Figure 2.**
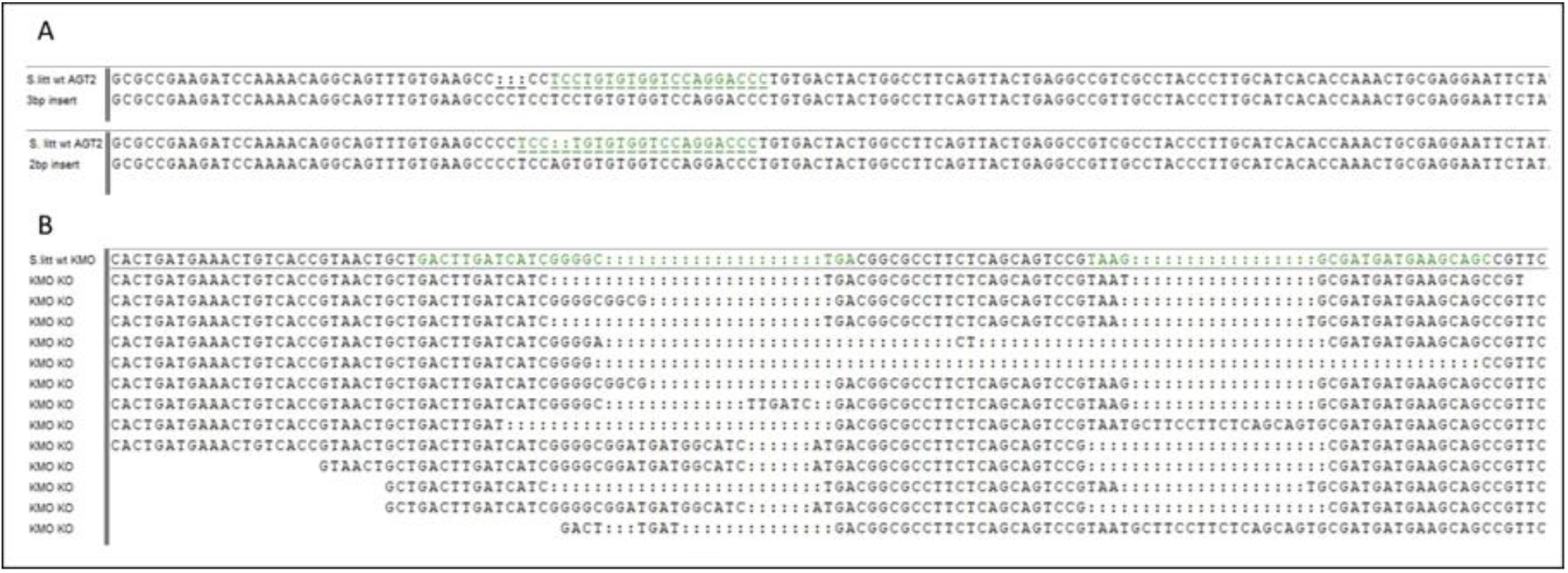
Mutated target sequences in *AGT2* and *KMO* genes. A) 3bp insert (above) and 2bp insert (below) occurred in the *AGT2* gene compared to the WT allele. The target sequence is highlighted in green in the WT. B) Examples of KMO-ko alleles compared to the WT allele. The two target sequences are highlighted in green in the WT.

Injections with *KMO* tracrRNA+crRNA resulted in many different mutant genotypes (Figure 2) and a visible phenotype in the adults. KMO-ko mutant adults showed bright golden eyes instead of dark brown eyes of the wild types (Supplementary Figure 1). Additionally, their wing and body scale color was lighter. This phenotype was subsequently used to identify homozygous KMO-ko mutants, regardless of their specific induced mutation and the line was maintained with a mix of mutant KMO-ko alleles.

### 2.3 Quantification of 8-HQA

Ten fifth instar larvae from each of KMO-ko (fed with 0 mM, 1 mM, 3mM, 5 mM and 10 mM 8-HQA) and WT conditions were starved for 5 hours and their regurgitant was collected in individual GC vials. To collect the regurgitant, each larva was pressed at the foregut area using a pair of forceps on a sterile petri dish, and the ejected fluid was collected using a pipette and weighed. The regurgitant samples were derivatized by means of tetrabutylammonium hydrogen sulfate (TBAS)-assisted anhydrous pentafluorobenzylation following a modified protocol published by (Naritsin et al., 1995), Figure 3a. This method involves addition of 250 μl of 0.1 M TBAS in 1M NH_4_OH solution at pH 10.4 to the regurgitant. After drying in a SpeedVac for 3 h, 400 μl of fresh pentafluorobenzyl bromide (PFBB)/N,N-diisopropylethylamine in acetonitrile solution (1.5 ml N,N-diisopropylethylamine, 0.6 ml PFBB and 27.9 ml dry acetonitrile, for 75 reactions), were added and heated at 65 ºC for 15 min without mixing. After cooling down to room temperature, 100 μL decane and 1 ml of sulfuric acid (1 M) were added and vortexed for 10 min at 1000 rpm. After centrifuging in a SpeedVac at 1,400 rpm (215 x g) for 10 min for phase separation the upper organic layer was transferred to GC-vials with 100 μL micro inserts.

**Figure 3.**
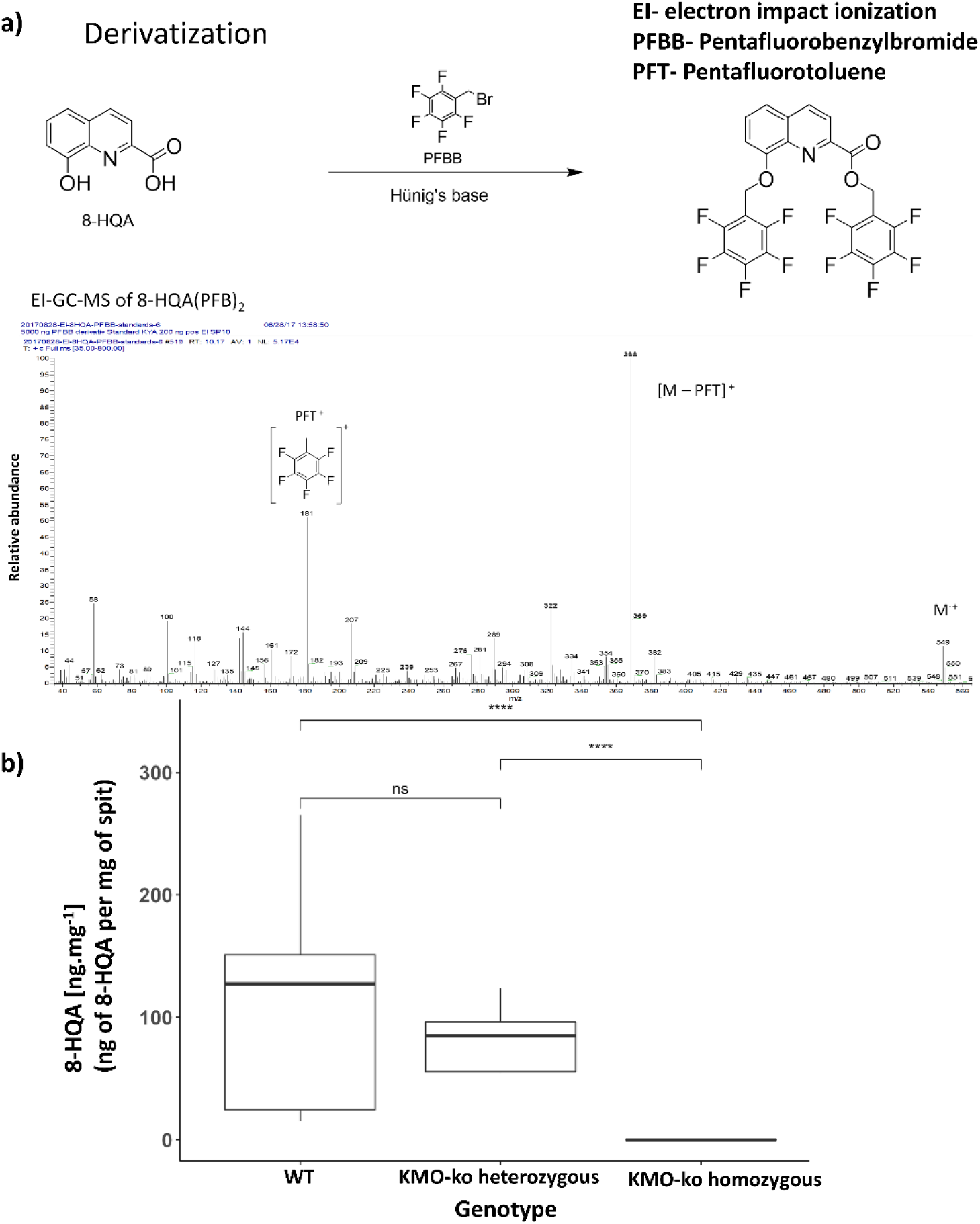
Detection and quantification of 8-HQA. a) Derivatization of 8-HQA with PFBB and mass spectrum of the derivatized compound. b) Concentration of 8-HQA in μg · ml^-1^ of regurgitant of the larvae in 5th instar wildtype (WT) and KMO gene knockout larvae. On the average, a WT larva has 142 μg · ml^-1^ of 8-HQA in the regurgitant (Wilcoxon test, p < 0.05).

Samples were analyzed by gas chromatography mass spectrometry (GC-MS) with an ITQ GC-ion-trap MS system (Thermo Scientific, Bremen, Germany) equipped with a fused silica capillary column ZB–5 (30 m × 0.25 mm × 0.25 μm with 10 m guard column, Zebron, Phenomenex, USA). Helium at 1 ml · min^-1^ served as carrier gas with an injector temperature of 320 °C running in splitless mode; 1 μL of sample was injected. Separation of the compounds was achieved under programmed temperature conditions (175°C for 1 min, then at 10 °C · min^-1^ to 320 °C kept for 1 min). The MS was run in EI mode (70 eV) with a scan range of 35 to 800 amu, a transfer line temperature of 320 °C, and an ion source temperature of 200 °C. Data acquisition was performed using Xcalibur 3.1 (Thermo Fisher Scientific). The basepeak m/z 368 at retention time 12.90 (corresponding the fragment [M-PFT]^+^) was used to plot a calibration curve and calculate the levels of 8-HQA in the samples.

### 2.4 DNA Extraction

DNA was extracted from 2nd, 3rd, 4th and 5th instar larvae using the DNeasy Power Soil kit (Qiagen). Prior to the extraction of DNA, each larva was homogenized with a bead mill (TissueLyser LT, Qiagen) for 3 minutes at a frequency of 29 Hz. The homogenate went through a Proteinase K treatment (50 μg · μL^-1^, Ambion) at 60 °C overnight. The extraction was done as per manufacturer’s protocol. The resulting DNA was purified using the Zymo Research DNA Clean & Concentrator-5 kit (Orange City, California). After the extraction and purification, the DNA concentrations of all the samples were measured using a NanoDrop One spectometer (Thermo Scientific) and stored at -20°C.

Each extracted sample DNA went through a quality control step which was a PCR amplification of the V4 region of the 16S rRNA gene. The PCR was conducted using forward primer 515F and reverse primer 806R (Table 2), The PCR was conducted using Thermo Fisher Scientific Taq DNA Polymerase (2 Units), 10 mM dNTP (0.2 mM), 50 mM MgCl_2_ (1.5 mM) and 0.5 μM of each forward and reverse primers. The PCR reaction cycle was an initial denaturation step at 94°C for 3 sec, followed 35 cycles of 94°C for 45 sec, 50°C for 30 sec, and a final elongation step at 72°C for 90 sec. Finally, an extended elongation step at 72°C for 10 min.

**Table 2:**
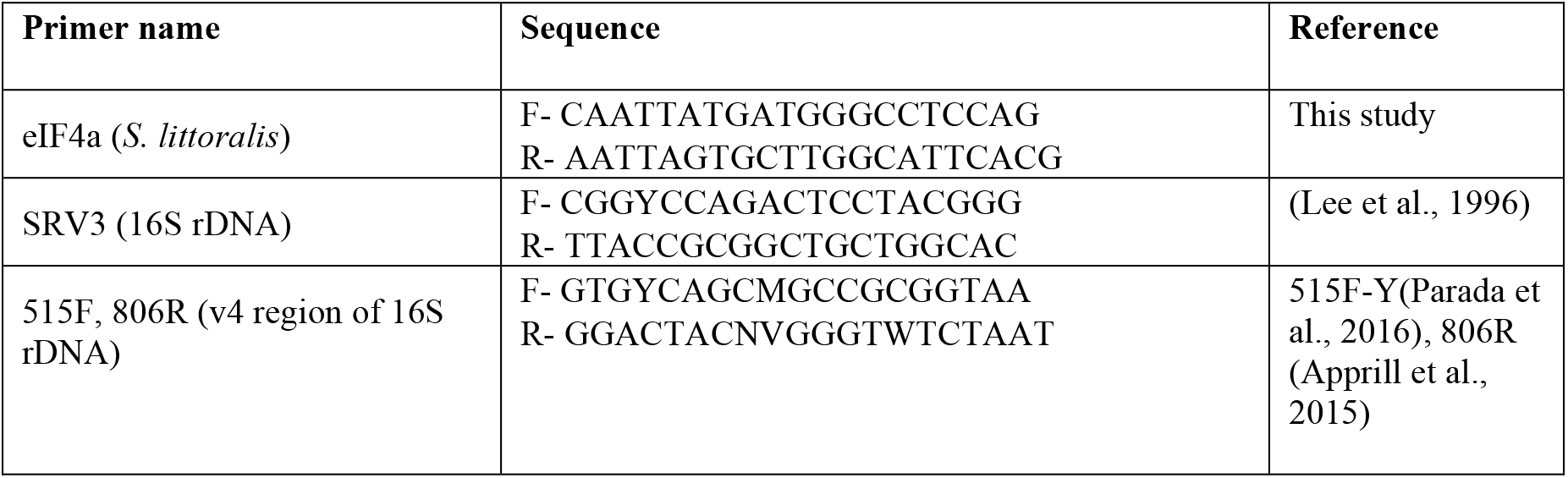
Primers used for quantitative PCRs and 16S amplicon sequencing.

The DNA marker that was used to determine the band size was Gene Ruler 1 kb Plus DNA Ladder. The samples were run on 1.5% Agarose gel containing ∼3-5 μl Midori green Advance DNA stain for visualization under UV illumination. The gel electrophoresis Bio-Rad Wide Mini-Sub® Cell GT chamber was used to run the samples at voltage of 150 V and 120 mA current for 35-40 min. The gel was visualized using Gel Doc™ XR+ System and band of ∼390 kb size was obtained.

### 2.5 Quantitative PCR Analysis

Each 10 μl PCR reaction consisted of 5 μl of SsoAdvanced Universal SYBR Green Supermix (Bio-Rad, Germany), 250 nM of forward and reverse primers: eIF4a (to amplify a host gene from midgut tissue) and SRV3 to amplify 16S bacterial rDNA (Lee et al., 1996) (Table 3), with the final volume made up with RT-PCR grade nuclease-free water (Thermo Fisher Scientific, Germany). The PCR reactions were conducted in the Bio-Rad CFX96 real-time PCR detection system. The reaction comprised the following steps: 98 °C for 3 minutes, followed by 35 cycles of heating at 98 °C for 15 seconds and 60 °C for 30 seconds. The melt curve was plotted from 65 to 95 °C in increments of 0.5°C. For each biological replicate, two technical replicates were measured. The C_t_ values were determined by log 10 fold-difference C_t_ = C_t_ (target gene)/C_t_ (reference gene), where the target gene is the 16S rDNA amplified by the SRV3 primers, and the reference gene is eukaryotic initiation factor 4a (eIF4a), a housekeeping gene of *S. littoralis*, used for normalization of the 16S rDNA copy numbers (Kang et al., 2019) relative to host DNA. The amplification efficiency of the two primer pairs was > 95%, and was determined with serial dilutions of the amplified gene products of SRV3 and eIF4a primers respectively.

**Table 3:** Anova (nested analysis of variance) table. Interaction was tested among genotype (KMO-ko/WT) of the Versailles larvae, the stage of the larvae (instar) and the technical replicates. The dependencies of log_10_ copy numbers (bacterial) to genotype (KMO-ko and WT) is highly significant (p= 2.37×10^−12^), and the fact that the interaction between genotype and instar is more significant (p= 0.0001) than the instar effect (p= 0.05) emphasizes the inconsistency of the genotype effect across instars. TukeyHSD post-hoc test showed significant differences in log_10_16S rDNA copy numbers between the KMO and WT genotypes of third, fourth and fifth instar stages.

|  | DF | Sum sq | Mean Sq | F value | P value |
| --- | --- | --- | --- | --- | --- |
| **Instar** | 3 | 9.79 | 3.26 | 4.767 | 0.005 |
| **Genotype** | 1 | 58.8 | 58.8 | 85.862 | $2.37 \times 10^{-12}$ |
| **Instar: Genotype** | 3 | 16.7 | 5.57 | 8.127 | 0.0001 |
| **Instar: Genotype: Replicate** | 8 | 0.47 | 0.06 | 0.086 | 0.99 |
| **Residuals** | 49 | 33.56 | 0.68 |  |  |

Standard curves to calculate the copy numbers were plotted by amplifying 16S rDNA (using SRV3 primers) from DNA of *Enterococcus mundtii* and eIF4a from surface-sterilized legs of *S. littoralis* adults. The PCR was conducted using Thermo Fisher Scientific Taq DNA Polymerase (2 Units), 10mM dNTP (0.2 mM), 50mM MgCl_2_ (1.5 mM) and 0.5 μM of each forward and reverse primer. The PCR reaction cycle was an initial denaturation step at 94°C for 3 sec, followed 35 cycles of 94°C for 45 sec, 60°C for 30 sec, and a final elongation step at 72°C for 90 sec. Finally, an extended elongation step at 72°C for 10 min. The segments of length 203 and 289 bp respectively were cut with a scalpel after running the amplified product in 1.5% agarose gel as described in S2. The cut gel segments were eluted using QIAquick Gel Extraction Kit (Qiagen). The extracted DNA was serially diluted (1 ng to 10^-8^ ng) and was used for plotting the respective standard curves.

To study the effect of larval stage and genotype on the bacterial numbers, the qPCR data were analyzed with the help of a nested ANOVA model as follows: the dependence of the ratio of gene copy number (copy numbers of 16S rDNA/eIF4a) was analyzed according to the stage of the insect, genotype (KMO-ko/WT), and additionally, the Ct value of technical replicates nested within the stage of the insect, which is further nested within the genotype. The model checks for the dependence on stage and genotype independently as well as their interaction. Tukey’s post-hoc test was done thereafter. The statistics were performed with R (version 3.6.3).

### 2.6 16S rRNA Gene Amplicon Sequencing and Analysis

Extracted DNA from 5 biological replicates of second, third, fourth and fifth instar larvae from KMO-ko and WT strains, and the KMO-ko strains fed with 0 mM, 1 mM, 3mM, 5 mM and 10 mM 8-HQA were sent for sequencing to MR DNA (Molecular Research LP), Texas. Sequencing of 2 × 250 bp paired-ends was performed on the IlluminaMiSeq platform using primer pair 515F (Parada et al., 2016), 806R (Apprill et al., 2015) (Table 2). Each extracted DNA sample went through a quality control step which was a PCR amplification of the V4 region of the 16S rRNA gene prior to sending for sequencing.

The QIIME 2.0 platform was used to analyze the sequencing data. Adapters and primers of the sequences were removed using the Cutadapt plugin (Estaki et al., 2020), (Martin, 2011) followed by denoising the sequences using the DADA 2 plugin (Estaki et al., 2020), (Callahan et al., 2016) (truncation of both forward and reverse reads was set at 230 and chimeras were removed using the consensus method). Classification of taxon was done by training Naive Bayes classifier using Greengenes reference sequences (Greengenes Database Consortium, version 13_8, clustered at 99% similarity) (DeSantis et al., 2006). Representative sequences from each OTU were generated and mapped against the Greengenes 16S database for taxon assignment. All samples were rarefied to the least number of reads (55599, 280187 pertaining to results 2 and 3 respectively). Taxa corresponding to mitochondria and chloroplast were filtered. R packages: Phyloseq and ggplot2 were used to visualize the analysis of the metagenomics data. Alpha diversity (Shannon index (Gauthier and Derome, 2021) and Faith PD indices) and Beta diversity were analysed by calculating the distances of Bray-Curtis (Beals, 1984) and Canberra (Warton et al., 2012) and were visualized as principal coordinate analyses (PCoA). The Adonis method for multivariate analysis of variance in the vegan package (Dixon, 2003) was used to check the variance among replicates of the different treatment groups. The Kruskal-Wallis test (P < 0.05) was carried out for each index of Alpha diversity, followed by the Wilcoxon test for pairwise comparison of KMO-ko and WT samples across developmental stages. R-packages-Phyloseq (McMurdie and Holmes, 2013), ggplot2 (Wickham, 2011), vegan (Dixon, 2003), ggpubr (Kassambara and Kassambara, 2020) were used for final visualization of data and statistics on R-studio using R version 3.6.3.

### 2.7 Data Availability

The raw genomic data have been deposited in the NCBI Short Read Archive (SRA). The BioProject ID is PRJNA727271. The *S. littoralis* genes have been deposited in GenBank (AGT1: GenBank Accession ON808443, AGT2: ON808444, KMO: ON808442). Additional data are in the Supplementary Information.

## 3 Results

### 3.1 8-HQA in KMO Knockouts and WT Larvae

In order to confirm the absence of 8-HQA in the larvae with mutants of the gene encoding kynurenine 3-monooxygenase, the concentration of 8-HQA was measured in the regurgitant of both WT and KMO-ko larvae. Figure 3b shows the levels of 8-HQA compared to the wild type (WT) counterparts (average 142 μg · ml^-1^of regurgitant). In KMO-ko individuals, 8-HQA could not be detected (Wilcoxon test p < 0.05) (Supplementary Data Sheet 1).

### 3.2 Microbial Diversity Across the Larval Instars of *S. littoralis*

The normalized 16S copy numbers (designating the bacterial abundance) in the KMO-ko are consistently higher than the corresponding WTs, along the stages of second, third and fourth instar (Fig. 2a). The nested ANOVA (Table 3) showed significant results (Tukey’s test, p < 0.05) for 3rd, 4th and 5th instars.

The KMO-ko and WT larvae showed differences in their bacterial composition over different developmental stages. Not only do KMO-ko harbor higher numbers of bacteria (Figure 4a) than the WTs, their bacterial composition is dominated to an even greater extent by the class of Bacilli of the phylum Firmicutes (Figure 4b, Supplementary Figure 3). The KMO-ko of the Versailles strain was fed with frass from the Syngenta strain because the previous experiments on the bacterial population of *S. littoralis* larvae were conducted on this strain (Tang et al., 2012;Teh et al., 2016). The frass mixed with the food itself had a dominance of Bacilli and the rest consisted of Actinobacteria and Proteobacteria (Supplementary Figure 2). Only in the 2nd instar KMO-ko can one find more diversity than the KMO-ko of other stages. This condition in particular harbours 5% and 2% of Beta- and Gammaproteobacteria, respectively. Although the Firmicutes are still the most abundant phylum, Actinobacteria, and Proteobacteria (Supplementary Figure 3) show their presence in differing levels among the WTs of the four larval stages. The 2nd instar shows a 10% and 3% population of Beta- and Gammaproteobacteria respectively, whereas the 4th instar harbors 16% Actinobacteria and 9% Gammaproteobacteria.

**Figure 4.**
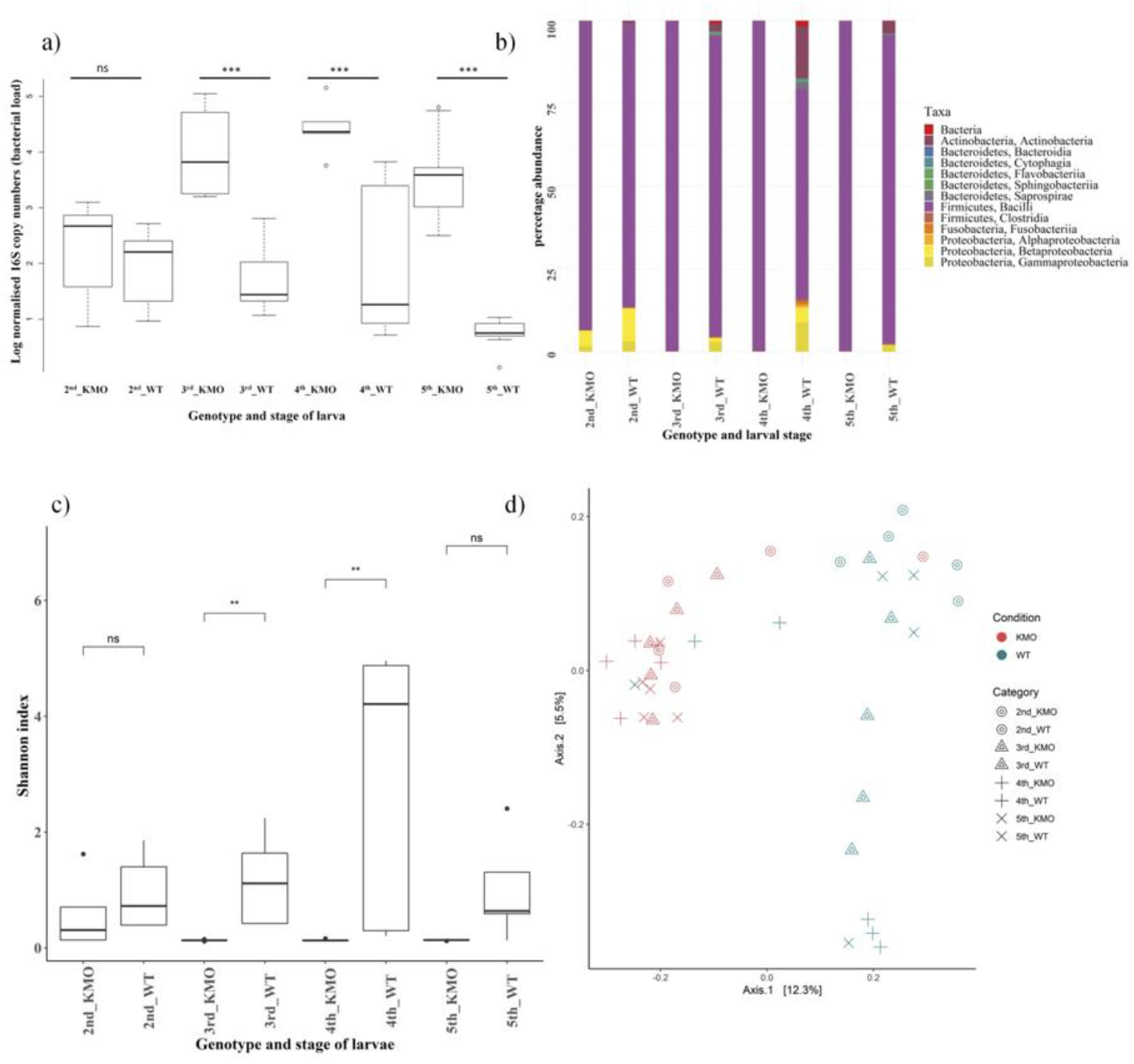
Bacterial abundance and composition in wild-type and KMO knockout larvae. a) 16S rRNA gene copy numbers signifying the bacterial abundance as determined by quantitative PCR during different stages of the life cycle of *S. littoralis* (2nd, 3rd, 4th and 5th instars) and the genotypes of KMO-ko and WT. The KMO-ko harbor a higher bacterial number than their corresponding WTs. b) Comparison of relative microbial composition at the phylum level in KMO-ko vs. WT *S. littoralis* through the larval instars. c) Box plot for comparison of alpha diversity of species in KMO-ko and WT guts of *S. littoralis*, through the larval instars, based on Shannon (species richness) index of alpha diversity (Wilcoxon test p < 0.05 for 3rd and 4th instars). d) Beta diversity analysis between the KMO-ko and WT lines of *S. littoralis* guts by Canberra distances on PCoA plot (Adonis test p=0.001). Each point represents an individual sample and clustering of points indicates similarity of bacterial composition in the individuals.

Alpha diversity was measured on the basis of Shannon (Figure 4c), and Faith phylogenetic diversity indices (Supplementary Figure 4). The Shannon index measures species diversity in terms of richness of distribution, while the Faith phylogenetic diversity index is based on phylogeny (Faith, 1992). The Kruskal-Wallis test (p < 0.05) followed by the Wilcoxon test was performed for pair-wise comparisons and it showed a significant difference (p < 0.05) between the two genotypes of KMO-ko and WT in the 3rd and 4th instars prominently in case of Shannon index (Figure 4c) (species richness), and 3rd instar when measured using Faith PD (Supplementary Figure 4).

Multivariate statistical analyses were employed to study the differences in bacterial composition across the genotypes (KMO-ko and WT) and individual stages (2nd, 3rd, 4th, 5th instars). Thus, beta diversity analysis was performed using Canberra distances (Warton et al., 2012) and Bray-Curtis (Beals, 1984) (Fig. 2d, S9), using Principal Coordinate analysis (PCoA) to analyze how the individual conditions cluster according to their taxonomic compositions. KMO-ko of all the four instars show clustering when the beta diversity is analysed using Canberra (Figure 4c) and Bray-Curtis distances (Supplementary Figure 5) on PCoA plots.

Analysis of variance (Adonis) using beta-diversity matrices-Bray-Curtis and Canberra distances gave significant results indicating that the bacterial composition was significantly different among the different conditions (p = 0.001) (Figure 4d, Supplementary Figure 5). Overlapping clusters especially among stages of WT or KMO-ko is due to several shared common taxa. Considering all the KMO-ko and WT together, they form two separate clusters on a PCoA plot according to their Bray-Curtis (Supplementary Figure 5) and Canberra distances (Figure 4d) because of their different taxonomic composition (Adonis, p=0.001).

### 3.3 Effect of Supplemental 8-HQA on KMO Knockout Larvae

KMO-ko larvae were fed with a range of concentrations of 8-HQA to study the effect on the microbial population. The WT condition was compared to KMO-ko on 0, 1, 3, 5 and 10 mM 8-HQA, added to their artificial diet. The 8-HQA recovered from the regurgitant was directly measured (Supplementary Figure 6, Supplementary Data Sheet 2). The concentration of 8-HQA from the KMO-ko larvae at 10 mM 8-HQA was closest to that of WT (Wilcoxon test, P > 0.5). In terms of bacterial content, these larvae were expected to behave more like WTs. Interestingly the bacterial numbers of 8-HQA-supplemented KMO-ko approached that of WTs (5th_WT) (ANOVA, P < 0.05, Tukey’s test between 0 mM 8-HQA and WT P < 0.05; whereas P > 0.05 for other 8-HQA concentrations vs. WT) (Figure 5a). However, the bacterial richness measured by the Shannon diversity index did not change in KMO-ko upon feeding with 8-HQA (Supplementary Figure 8, Wilcoxon test p < 0.05 for 8-HQA-fed KMO-ko vs. WT). They harboured 99% Enterococci (Figure 5b, Supplementary Figure 7), similar to the control KMO-ko (0 mM 8-HQA and 5th_KMO), as opposed to 64% Enterococci and 30% Betaproteobacteria in the WTs (5th_WT). The beta diversity analysis using Canberra distances show clustering of 5th instar WTs away from all the KMO-ko fed with 8-HQA, suggesting that an external administration of 8-HQA to the KMO-ko is not recapitulating the bacterial composition in KMO-ko as that of WTs (Supplementary Figure 9).

**Figure 5.**
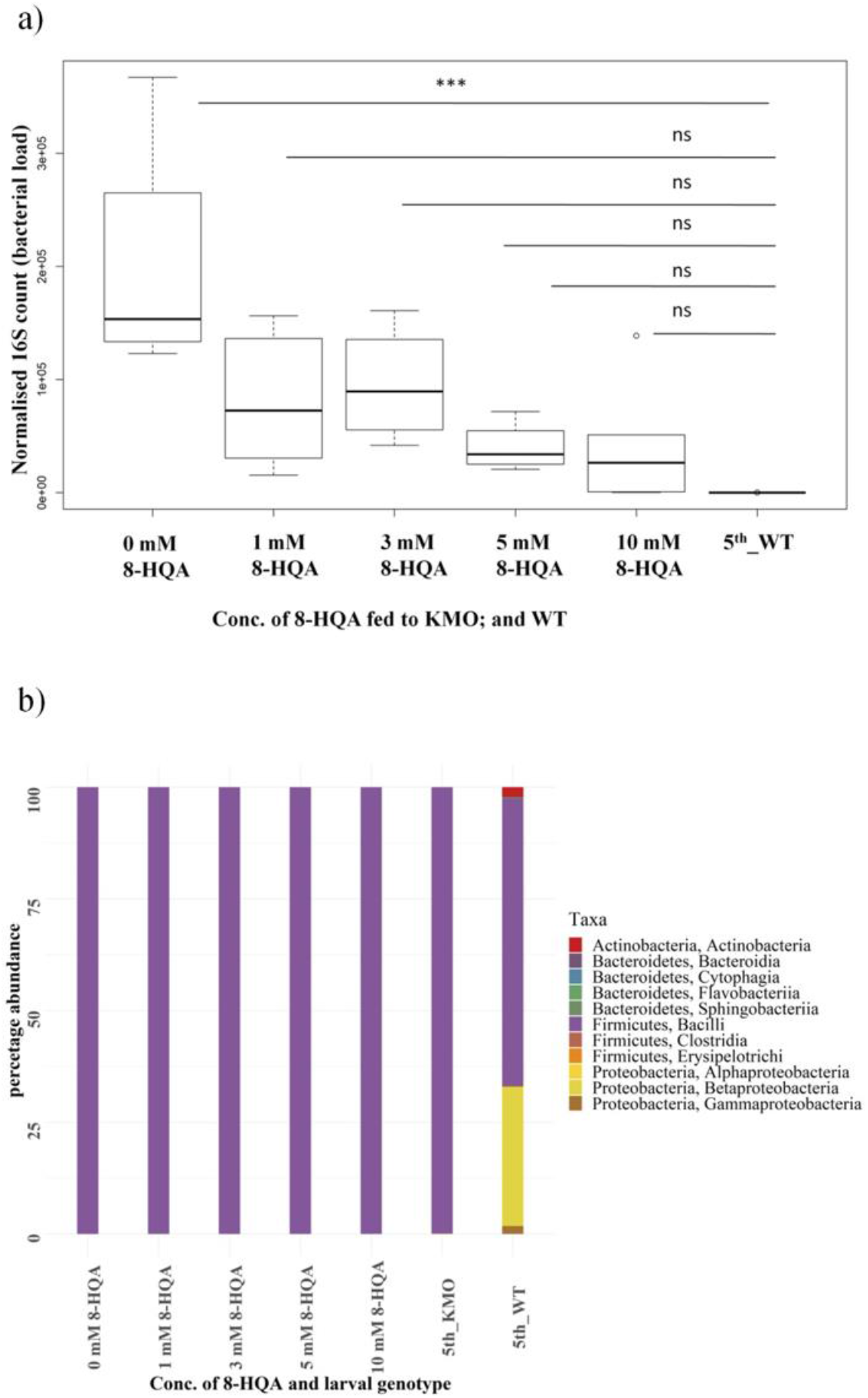
Results of feeding 8-HQA to KMO knockout larvae. a) Normalized 16S copy numbers of the same groups of larvae, a measure of the bacterial abundance in each condition: 5th instar larvae-KMO-ko fed with 0, 1, 3, 5, 10 mM 8-HQA, and WT. The bacterial numbers of KMO-ko approach that of WT upon administration of the compound (ANOVA, p < 0.05, Tukey’s test between 0 mM 8-HQA and WT < 0.05). b) Bacterial composition of KMO-ko larvae fed with the same increasing concentration of 8-HQA, KMO-ko control, and WT. The taxa distribution of KMO-ko do not resemble that of WTs, and Bacilli is the most abundant class in KMO-ko even after administration of 8-HQA.

## 4 Discussion

Previous studies (Tang et al., 2012;Pesek et al., 2015;Teh et al., 2016) have confirmed the presence of a high concentration of 8-HQA in the guts of *S. littoralis* but none of them studied the impact of the compound on the microbiota of the host. To test how the gut microbial diversity is influenced by the presence of the compound 8-HQA, the gene encoding kynurenine 3-monooxygenase (KMO) from the biosynthetic pathway (Figure 1) was knocked out in a strain of *S. littoralis*. The microbial diversity of these strains was compared to that of wild type strains (WT) that produced normal levels of the compound.

KMO-ko had a higher number of bacteria than WT, while WT had higher species richness--the number of different species represented in each condition as measured by the Shannon alpha diversity index. Most of the reads assigned to KMO-ko hit the OTU of Firmicutes (family Enterococcaceae) (Figure 4b, Supplementary Figure 2), which leaves negligible room for other phyla. While the 2nd instar KMO-ko contain some levels of Alphaproteobacteria, their bacterial richness is still less than their corresponding WTs. Thus, in spite of seemingly housing a greater number of bacteria in the gut, the KMO-ko knockouts of *S. littoralis* show a lower diversity than the corresponding wild types. Administration of external 8-HQA to the KMO-ko to mimic the gut concentration of WTs only restored the bacterial numbers, but not the diversity.

In the gut of *S. littoralis* larvae, the sequestering ability of 8-HQA for both Fe^2+^ and Fe^3+^ is pH dependent. At an acidic pH, it is greater for Fe^3+^, whereas at an alkaline pH, it is higher for Fe^2+^. The alkaline foregut therefore has more ferrous ions bound to the compound, leaving the ferric ions to form hydroxo species. In the neutral hindgut, the complex of 8-HQA is more prone to form. Also, the binding affinity of 8-HQA to these cations is so high and stable, that it leaves hardly any free cations to hydrolyze (to ferrous and ferric hydroxides) in the environment of high concentration of 8-HQA (Gama et al., 2018). In KMO-ko however, in absence of the iron chelator 8-HQA, presence of the hydrolyzed species is expected. We expect certain metabolic changes to occur when iron is chelated vs. when it is available in abundance. Iron dependent enzymes are a key to energy acquisition pathways. Change in iron levels could change the redox balance. In a similar study, acetate that resulted from a fermentative pathway was reduced, and at the same time there was a reduction in fermentative bacteria (Parmanand et al., 2019).

A study on broiler chickens with and without iron fortified diet showed similar results as ours. The bacterial diversity in the guts of chicken underfed on iron showed a higher bacterial diversity than the ones with excess iron in their diet (Reed et al., 2017). So was the case in rhinoceros prone to iron overload disease (Roth et al., 2019). In such studies the researchers hypothesized that a control over the iron levels could keep levels of pathogens in check (Parmanand et al., 2019). Although we did not find any visible health deficits in the KMO-ko knockouts, earlier studies have shown the pathogenicity of the larva’s own Enterococcal species-*E. fecalis* and *E. fecium* (Tang et al., 2012). It has been established that *Enterococcus mundtii* depletes the levels of the opportunistic pathogens by means of its bacteriocin, called mundticin (Shao et al., 2017), yet we still do not know if 8-HQA is playing an additional role in keeping these pathogens under control. In a study with another Lepidopteran, *Galleria mellonella*, researchers found that both the host lysozyme and mundticiin produced by its symbiont, *E. mundtii* are acting synergistically to keep pathogens in check (Johnston and Rolff, 2015). In our case as well, it could be quite possible that the host factor like 8-HQA is controlling the levels of the Enterococci, since the opportunistic pathogens belong to this genus. 8-HQA is possibly doing so by allowing Proteobacteria to grow in addition to simultaneously increase the gut bacterial diversity, and by reducing the overall bacterial numbers. Since we lack species-level resolution, we are unaware of the exact species of the Enterococcal population that are present in the KMO-ko knockouts vs. the WT.

The community of gut bacteria in *S. littoralis* is not just affected by 8-HQA, but also by the food consumed by the larvae. In this case, both the WT and KMO-ko larvae have been fed with the same artificial diet inoculated with frass from the Syngenta strain, and their colonization was monitored in the two genotypes. Needless to say, keeping all the dietary factors constant, solely administering 8-HQA will not introduce any new bacteria in the KMO-ko. It seems that in absence of 8-HQA, Firmicutes outgrow Proteobacteria inside the host gut, which remains unchanged upon administering 8-HQA to the KMO-ko. However, higher bacterial densities indicate that at least cell numbers could be rescued as compared to WT, which is in support of our hypothesis.

## 5 Acknowledgements

We thank Angelika Berg and for rearing insects and Dr. Nico Ueberschaar (MS-Platform, Friedrich Schiller University in Jena) for use of the ITQ for the final GC-MS measurements.

## 6 Contribution to the Field Statement

The microbial community of the digestive system of herbivorous caterpillars is determined by many factors. Intake from the food source may be highly variable. Competitive interactions among different bacterial species play a role. Regulation by the host’s immune system may limit and shape species composition, yet manipulation of the host’s means of control is rarely performed. We investigated how the host may control microbial diversity and abundance by restricting access to iron, a limiting nutrient. Many noctuid caterpillars, including the cotton leaf worm, secrete a potent iron chelator, 8-hydroxyquinoline-2-carboxylic acid, into the midgut. We abolished production of this chelator by knocking out a biosynthetic enzyme in the insect with CRISPR/Cas9. Significant effects on bacterial abundance and a minor influence on diversity resulted, showing that non-immune mechanisms may also play a role in host-microbiome interactions.

## 8 Author Contributions

TM planned the study, isolated insect and bacterial DNA, prepared samples for sequencing, analyzed data, and wrote the manuscript. SH generated and screened the knockout insect lines. SPS analyzed the bacterial sequence data, assisted with qPCR and conducted statistical analysis. AM standardized methods of DNA extraction and 16S rDNA amplification. SB set up derivatization, GC-MS measurements and quantification of 8-HQA. DGH planned the study and revised the manuscript. WB directed the overall project, planned the study, and revised the manuscript.

