## Supplementary material for "8-HQA Adjusts the Number and Diversity of Bacteria in the Gut Microbiome of *Spodoptera littoralis*"

**Supplementary Figure 1.  Adult eye color for (A) WT  and (B) KMO-ko**

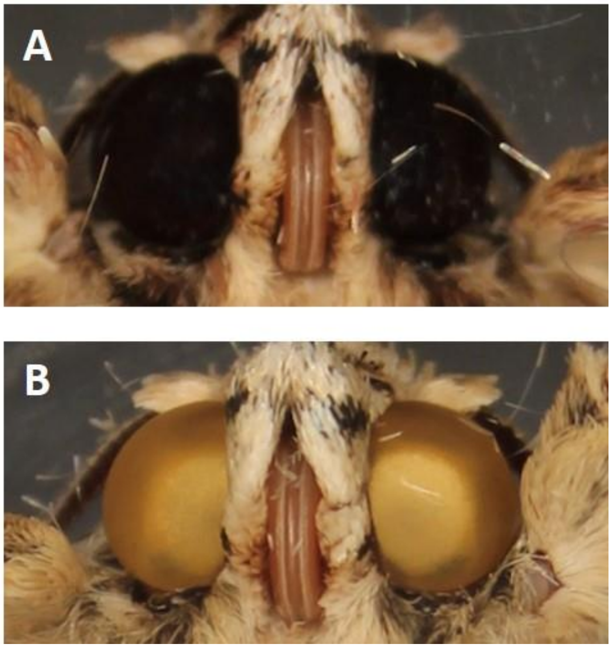

**Supplementary Figure 2.** Relative microbial composition of the frass of *S. littoralis* of Syngenta strain in artificial diet that was fed to the Versailles larvae because the previous experiments (Teh, Apel, Shao, & Boland, 2016) on the bacterial population of *S. littoralis* larvae were conducted on this strain.

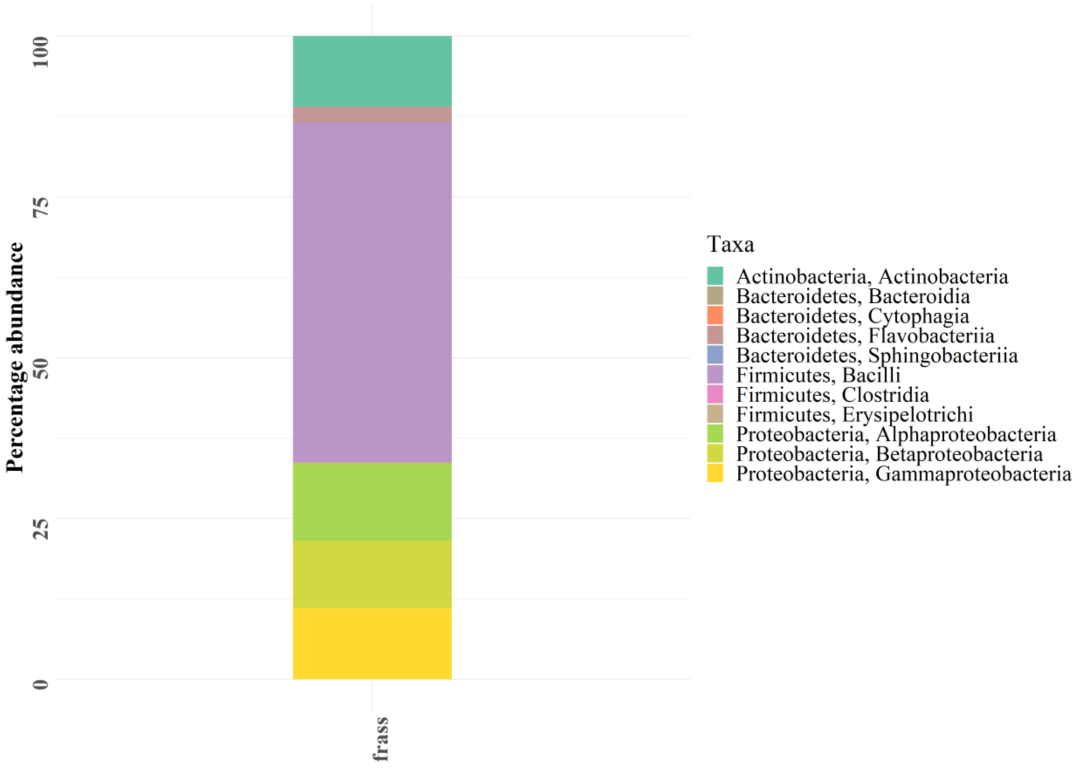

**Supplementary Figure 3**. Taxonomic composition of the two most dominant taxa: a) Firmicutes and b) Proteobacteria along the stages of life cycle of *Spodoptera littoralis* in their KMO-knock-out and WT genotypes (Wilcoxon test, $p < 0.05$ at the significant differences marked).

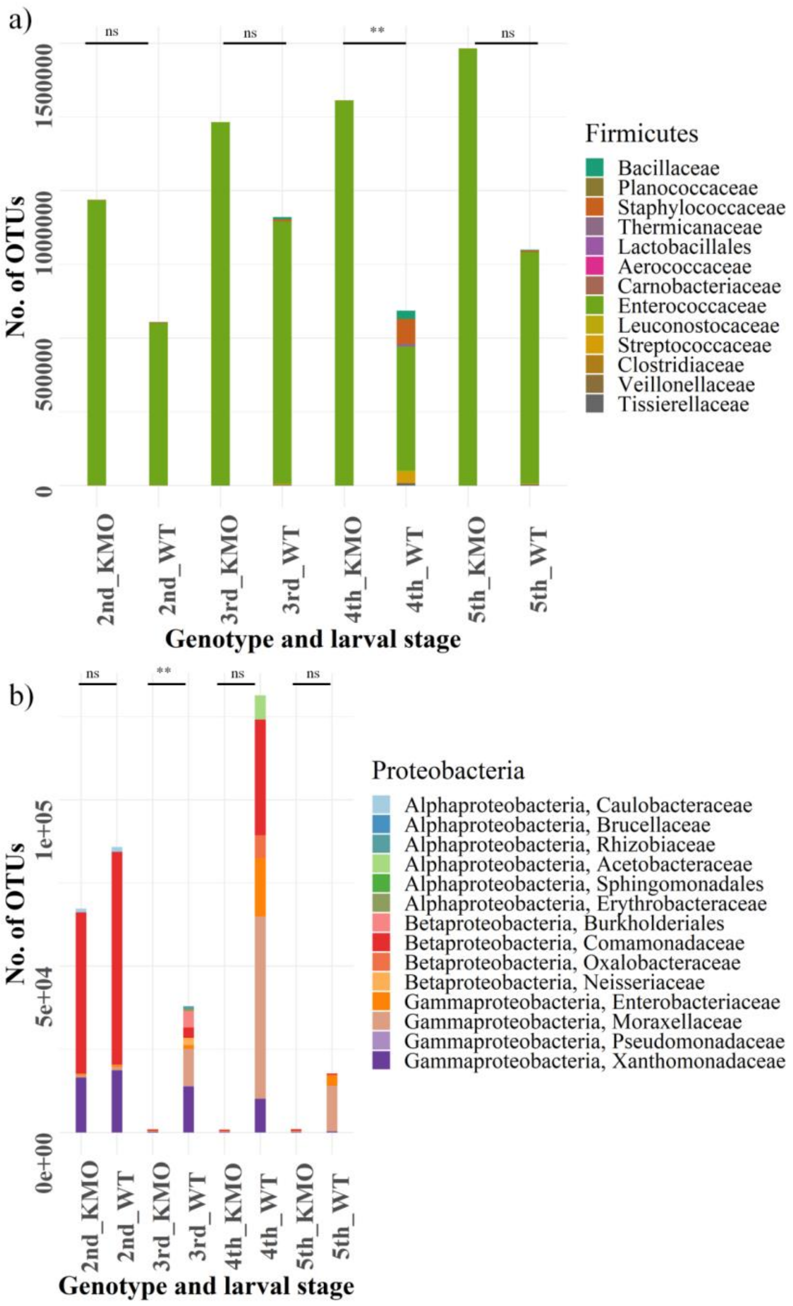

**Supplementary Figure 4.** Box plots for comparison of alpha diversity of species in KMO and WT guts of *S. littoralis*, along their life cycle (second, third, fourth, fifth instars), based on Faith-phylogenetic index (Faith, 1992) of alpha diversity (Kruskal-Wallis test $p < 0.05$ followed by pair-wise Wilcoxon test – $p < 0.05$ between 3rd_KMO and 3rd_WT).

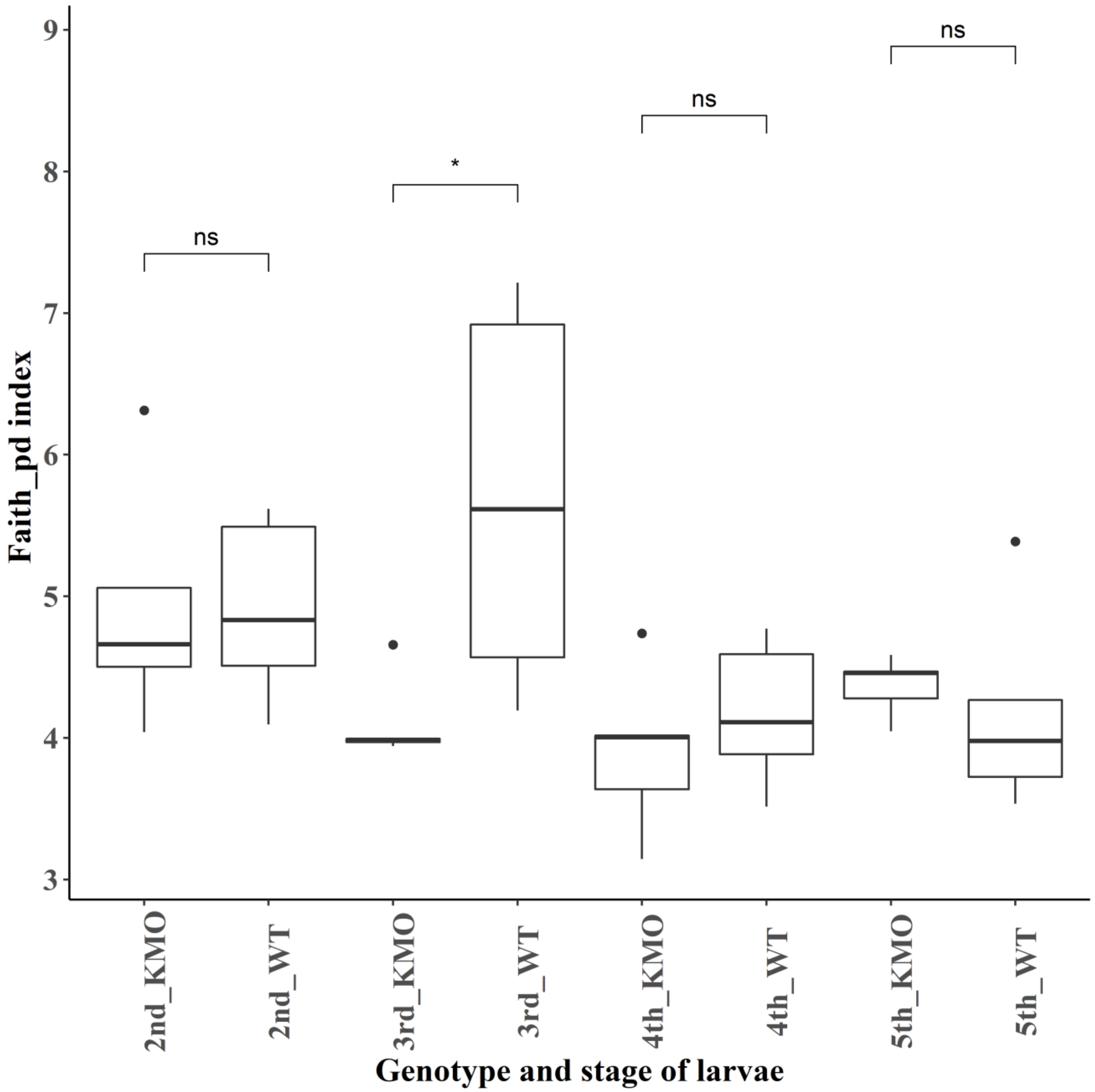

**Supplementary Figure 5.** Beta diversity analysis between the KMO and WT lines of *S. littoralis* guts by Bray-Curtis distances on PCoA plot (Adonis test p=0.001). Each point represent individual samples and clustering of points indicate similarity of bacterial composition in the individuals. The bacterial composition of the knock-outs are different from that of WTs.

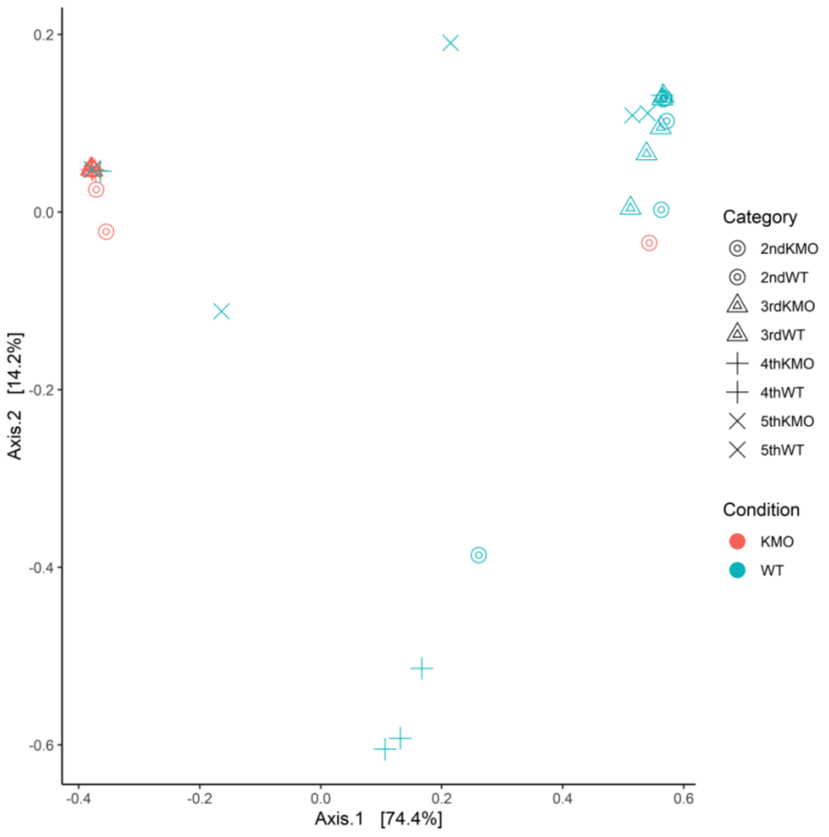

**Supplementary Figure 6.** Concentration of retrieved 8-HQA from the regurgitant of the KMO-ko larvae fed with an increasing concentration of 8-HQA (Wilcox test between KMO-ko and WT; and 1 mM and WT, $p < 0.05$). When the knockout larvae are administered with 10 mM of 8-HQA, the concentration of the compound retrieved in their regurgitate is the most similar to the regurgitant of the WT (supplementary sheet S15).

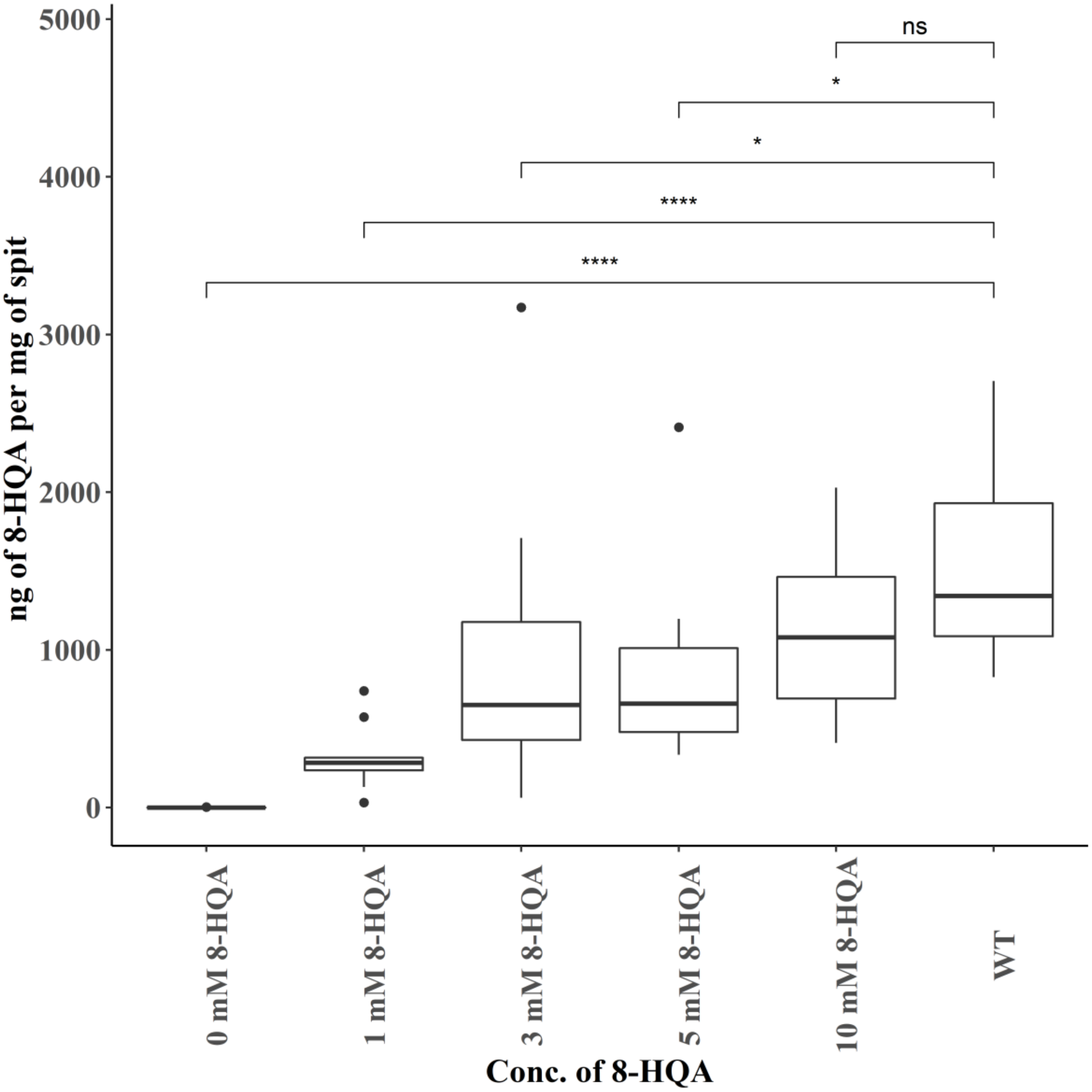

**Supplementary Figure 7.** Bacterial composition of the most dominant taxon of Firmicutes when KMO-ko larvae were fed with an increasing concentration of 8-HQA, and WTs. The WT condition consists of the least numbers of Firmicutes than the knockouts, or the knockouts administered with external 8-HQA (Wilcoxon test, $p < 0.05$, in 0 mM, 1 mM, 3 mM vs. 5th_WT).

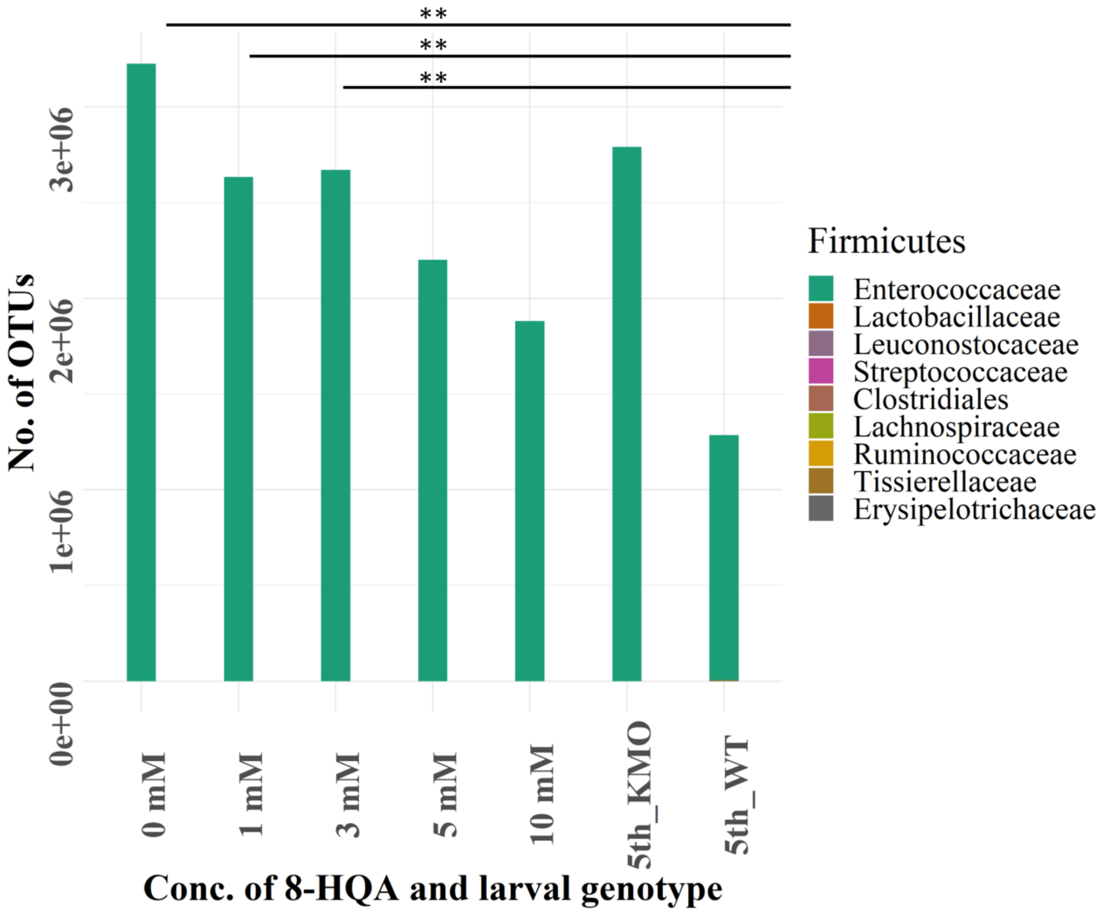

**Supplementary Figure 8.** Box plots for comparison of alpha diversity of 5th instar KMO-knock-out *S. littoralis* larvae fed with an increasing concentration of 8-HQA (0, 1, 3, 5, 10 mM); KMO control and WTs, measured using Shannon diversity index (Kruskal-Wallis test $p < 0.05$ followed by pair-wise Wilcoxon test, $p < 0.05$).

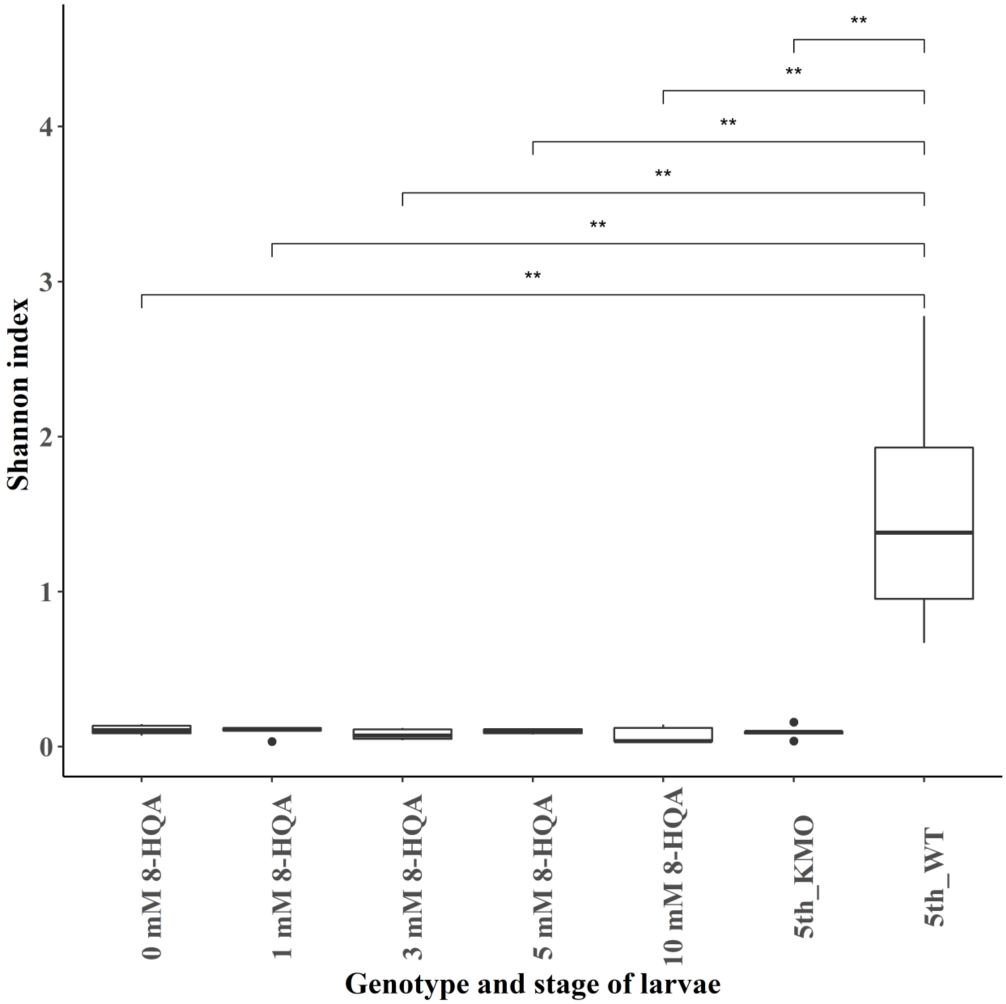

**Supplementary Figure 9**. Beta diversity analysis measured using Canberra distances (on a PCoA plot) showing that the bacterial composition of 5th instar WT is different from the KMO-ko with added 8-HQA (Adonis test $p < 0.05$). Each point represents individual samples and clustering of points indicate similarity of bacterial composition in the individuals. The bacterial composition of the 5th instar WT (5thWT) is different from all the other conditions, and feeding of an increasing concentration of 8-HQA to KMO-knockouts does not restore their bacterial composition to that of WTs.

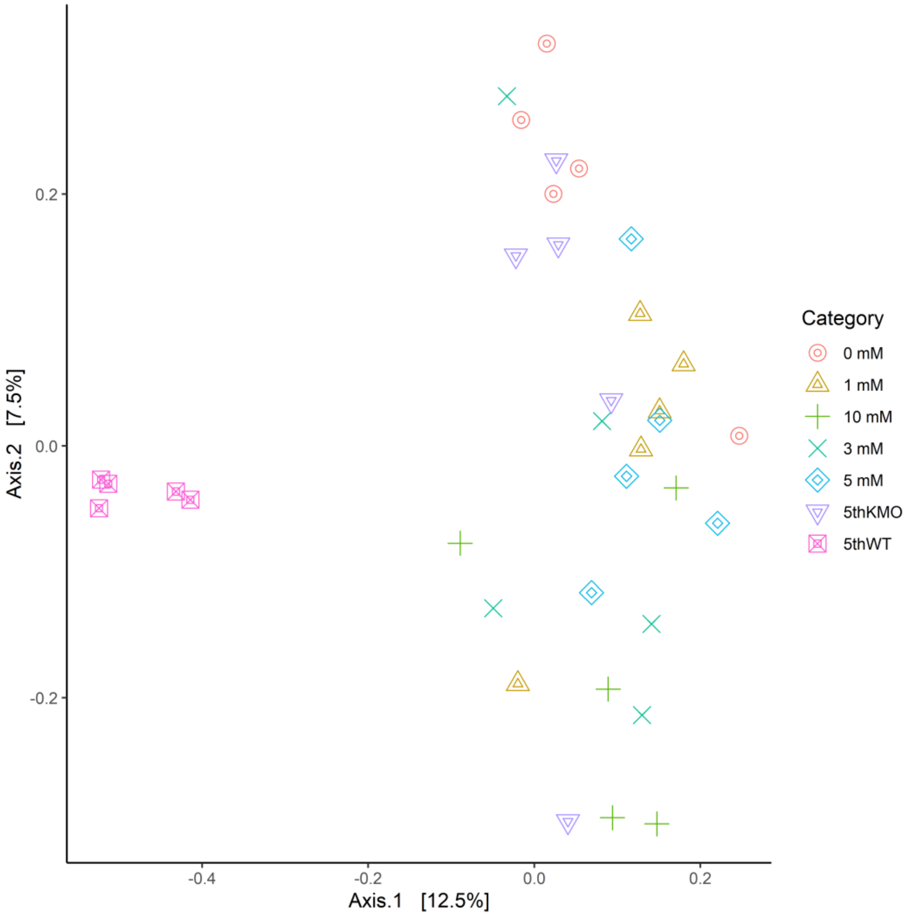

Supplementary sheet S14- GC-MS measurements of 8-HQA in KMO-ko and WT *S. littoralis* larval regurgitant, at 5th instars.

Supplementary sheet S15- GC-MS measurements of 8-HQA in KMO-ko *S. littoralis* larval regurgitant, at 5th instars that were fed with and increasing concentration of 8-HQA

**References to Supplementary Material**

Faith, D. P. (1992). Conservation evaluation and phylogenetic diversity. Biological Conservation, 61(1), 1-10.

Teh, B.-S., Apel, J., Shao, Y., & Boland, W. (2016). Colonization of the intestinal tract of the polyphagous pest *Spodoptera littoralis* with the GFP-tagged indigenous gut bacterium *Enterococcus mundtii*. Frontiers in Microbiology, 7, art. 928.
